# FveDAD2 negatively regulates branch crowns by affecting abscisic acid metabolism through FveHB7 in woodland strawberry

**DOI:** 10.1101/2024.01.02.573951

**Authors:** Hongying Sun, Junxiang Zhang, Weijia Li, Yan Wang, Zhihong Zhang

**Affiliations:** Liaoning Key Laboratory of Strawberry Breeding and Cultivation, College of Horticulture, Shenyang Agricultural University, Shenyang, 110866, China; Engineering Research Center of Coal-Based Ecological Carbon Sequestration Technology of the Ministry of Education, Shanxi Datong University, Datong, 037009, China

## Abstract

Strigolactones (SLs) are a significant hormone in plant growth response, crucial for regulating branching. DECREASED APICAL DOMINANCE2 (DAD2) is a novel receptor of SL. Here, FveDAD2 in woodland strawberries (*Fragaria vesca*) as the receptor for SL was identified, and three FveDAD2-RNAi transgenic lines that exhibited the phenotype of multi-branched crowns and smaller fruits were obtained. Gene expression, phenotypic analysis, and yeast assays were used to investigate the function of FveDAD2 in regulating branch crowns in strawberries. Like the alpha/beta hydrolase D14, FveDAD2 interacts with FveSMXL7 and depends on SL. Furthermore, the yeast single-hybrid, GUS activity assay, and LUC assay results demonstrate that FveSMXL7 binds to the promoter of *FveHB7* and repress its transcription. FveHB7, a homeobox transcription factor, negatively regulates the transcription of *FveABA8’OH1*, which encodes the enzyme that catabolizes abscisic acid (ABA). ABA contents were reduced in the shoot tips of the FveDAD2-RNAi lines, while treating wide-type plants with 20 μM ABA significantly suppressed the number of branches. In conclusion, we discovered a novel pathway of SL signaling to regulate branching through ABA.

**One-sentence summary:** FveDAD2 negatively regulates branch crowns by affecting abscisic acid levels by interacting with FveSMXL7 to regulate the expression of *FveABA8’OH1* via the transcription factor FveHB7.

## Introduction

Shoot branching is a crucial agronomic trait that plays a vital role in plant development, which significantly shapes plant architecture to achieve increased yield potential across various plant species (Kelly et al., 2023). Bud growth is regulated by various endogenous and environmental signals, including the phytohormone strigolactones (SLs). SLs were initially recognized as significant inter-root signaling molecules produced by plants that served as stimulants for the germination of parasitic plants and proved to enhance clumped mycorrhizal symbiosis. SLs are a group of compounds derived from carotenoids that are critical to inhibit branching (Gomez-Roldan et al., 2008; Umehara et al., 2015; Chesterfield et al., 2020). Much has been learned about the biosynthesis and signaling of SLs by identifying and examining branching mutants in some species. The SL biosynthetic pathway seems to be conserved, but regulating the shoot branching by the SL signaling pathway is still undetermined in higher plants.

The perception of SLs involves α/β hydrolase-(DWARF14) D14, RAMOSUS3 (RMS3), and DECREASED APICAL DOMINANCE2 (DAD2). Once the receptor hydrolyzes SL, the SL is divided into the ABC-ring and D-ring (Simons et al., 2007; Hamiaux et al., 2012; de Saint Germain et al., 2016; Yao et al., 2016; Yao et al., 2018). Then, the receptor covalently captures the D-ring and forms an SCF complex with the MAX2/D3/RMS4 F-box and D53/SUPPRESSOR OF MAX2-LIKE 6,7,8 (SMXL6,7,8) proteins, which leads to ubiquitination-dependent degradation of D53/SMXL6,7,8 (Zhou et al., 2013; Liang et al., 2016; Shabek et al., 2018; Kerr et al., 2021). D53 interacts with TOPLESS (TPL) and TOPLESS-RELATED (TPR) as well as specific SQUAMOSA PROMOTER-BINDING PROTEIN-LIKE (SPL) proteins, to repress the transcription of SPLs (Smith and Li, 2014; Soundappan et al., 2015; Wang et al., 2015; Liu et al., 2017; Ma et al., 2017; Song et al., 2017; Xie et al., 2020; Sun et al., 2021). Additionally, SMXL6,7,8 can directly bind DNA and repress the expression of *SMXL7* in *Arabidopsis*, acting as autoregulated transcription factors (TFs) to ensure SMXL6,7,8 homeostasis and negatively regulate SL signaling. (Wang et al., 2020).

Multiple factors, including hormones, nutrients, and sugars, regulate the branching of the SL pathway. SMXL6,7,8 is crucial for the influence of SLs on the accumulation of PIN-FORMED (PIN) proteins and the transport of auxin, which controls bud establishment (Crawford et al., 2010; Scheres et al., 2013; Soundappan et al., 2015; van Rongen et al., 2019; Zhang et al., 2020). SLs can potentially regulate cytokinin biosynthesis and metabolism via D53/SMXL6,7,8 (Duan et al., 2019; Cao et al., 2023). Additionally, cytokinins upregulate shoots’ SMXL7/D53 expression (Kerr et al., 2021). SLs also affect meristem formation via D53 at different nitrogen concentrations by changing the binding of the gibberellin negative regulator SLENDER RICE1 (SLR1) to the TF GROWTH-REGULATING factor 4 (GRF4) (Sun et al., 2023). The downstream factor regulating shoot growth of the SL repressor D53 is *TEOSINTE BRANCHED1* (*TB1*)/*FINE CULM1* (*FCL1*)/*BRANCHED1* (*BRC1*), which are highly expressed in axillary buds and inhibit bud growth (Aguilar-Martínez et al., 2007; Braun et al., 2012; Wang et al., 2015; Liu et al., 2017; Wang et al., 2020). The TB1/BRC1 pathway is linked to ABA and is important in controlling branching. Under short photoperiods, BRC1 induces *HOMEOBOX 21* (*HB21*), *HB40*, and *HB53* expression. These TFs with BRC1 further stimulate the *9-CISEPOXICAROTENOID DIOXIGENASE 3* (*NCED3*), leading to the accumulation of ABA in shoots, which affects shoot growth (González-Grandío et al., 2016; Dong et al., 2019). The TF NITROGEN-MEDIATED TILLER GROWTH RESPONSE 5 (NGR5) is involved in nitrogen-promoted tillering and is regulated through *D14* (Wu et al., 2020). Sugar is an initial regulator of shoot growth in SL signaling by regulating the BRC1 and cytokinins (Mason et al., 2014; Fichtner et al., 2017; Bertheloot et al., 2019; Barbier et al., 2021; Patil et al., 2021; Salam et al., 2021; Wang et al., 2021). In addition, the regulator CIRCULAR CLOCK ASSOCIATED1 (CCA1) can regulate tillering by modulating the SL signaling (Wang et al., 2020).

Regulating branching through signaling pathways with SLs is challenging, and the existing research results are difficult to integrate into a generalized model. Here, we obtained FveDAD2-RNAi transgenic lines of woodland strawberries (*Fragaria vesca*) with more branch crowns than wild-type plants. We found that ABA levels were significantly decreased at the shoot tips of multibranched FveDAD2-RNAi plants. Interestingly, we identified that FveDAD2 affects ABA levels by interacting with FveSMXL7, which indirectly regulates the expression of *FveABA8’OH1* in the ABA metabolic pathway via the TF FveHB7. These results indicate a novel pathway for ABA involved in the SL signaling pathway to regulate branch crowns.

## Results

### FveDAD2 is a conserved α/β-hydrolases superfamily protein

To investigate how SL regulates branch crowns in the woodland strawberry, the amino acid sequence of AtD14 was used to search for homologs in the Genome Database for Rosaceae (GDR). A homolog protein of woodland strawberry with 269 amino acids showed 76.58% similarity with AtD14 (Supplemental Fig. S1A) was identified. The annotation of this protein in the National Center for Biotechnology Information (NCBI) is strigolactone esterase DAD2 (*Fragaria vesca*), so we named it FveDAD2.

To establish the evolutionary relationship between FveDAD2 and other homolog proteins, 44 sequences from various species were obtained to conduct phylogenetic analysis. We found FveDAD2 is closely related to RcDAD2 (Fig. 1A). Using the MEME online website (https://meme-suite.org/meme/doc/meme.html), we analyzed the conserved structural domains of DAD2 in 10 different species. We found that there are eight motifs of α/β helical structures in FveDAD2, which corresponds to these proteins of dicots, and the N-terminal of monocot proteins have an additional glycine- and serine-rich (Fig. 1B). The putative catalytic triad (serine, aspartate and histidine) is necessary for SL hydrolysis, which is conserved among the three sites (Supplemental Fig. S1B FveDAD2: Ser98, Asp199, His248). Additionally, four putative crucial amino acids are used to identify components of the SCF complex (Supplemental Fig. S1B FveDAD2: Gly160, Pro163, Glu176, Ser178; green arrow). Taken as a whole, these data indicate that DAD2 was highly conserved in different plant species.

**Figure 1.**
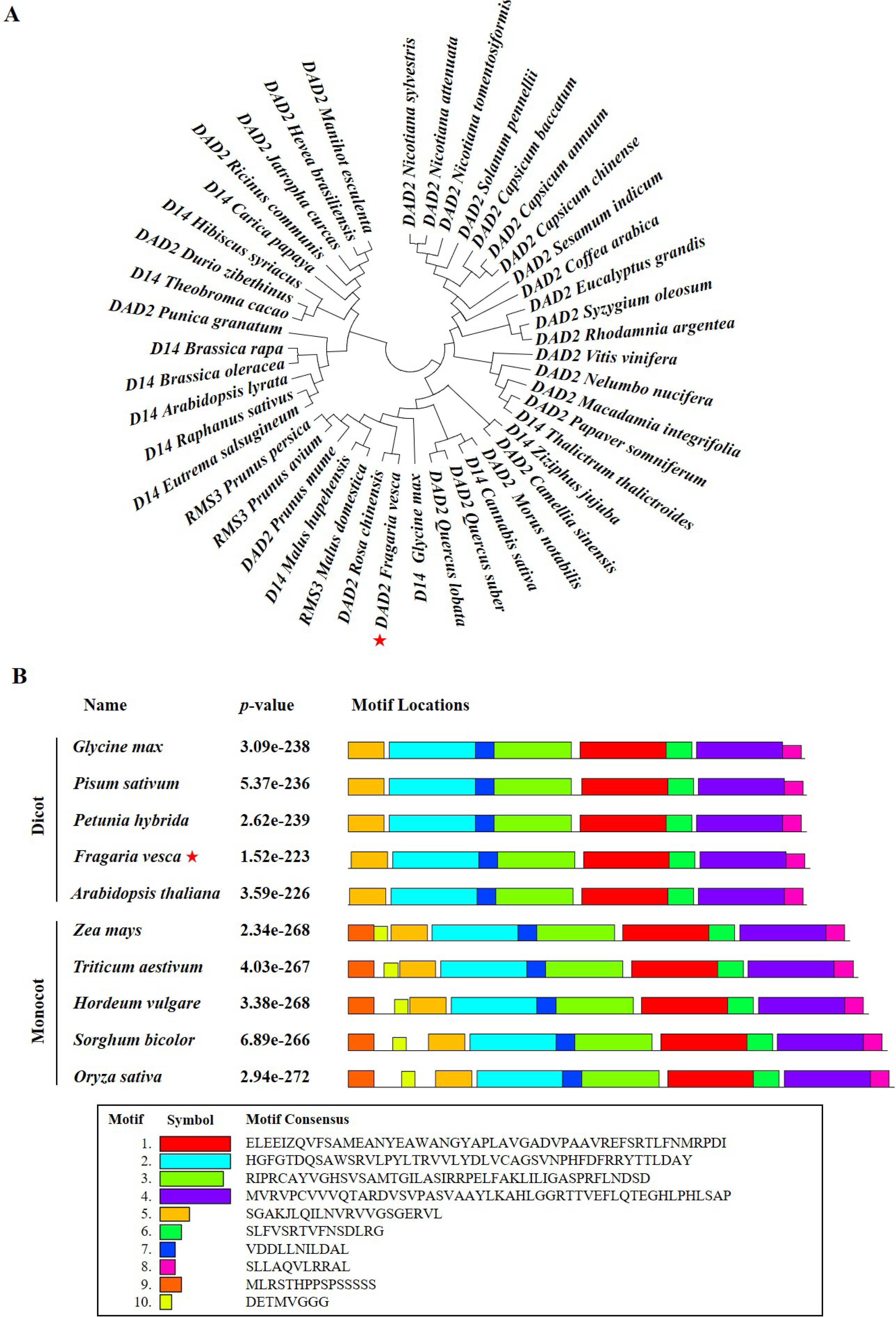
Identification and phylogenetic analysis of FveDAD2 **A)** Phylogenetic tree of FveDAD2 using MEGA6 by the Neighbor-Joining method. Specific sequence information is shown in Table S1. **B)** Conservative domain analysis of FveDAD2 and homologous proteins from the monocots and the dicots using MEME.

## *FveDAD2* is localized in the nucleus and expressed in different organs

To analyze the subcellular localization of FveDAD2, the pro35S::GFP constructs and pro35S::FveDAD2-GFP were transiently expressed in the leaves of *Nicotiana benthamiana*. It showed that the pro35S::FveDAD2-GFP fusion protein was localized in the nucleus. In contrast, the pro35S::GFP protein was detected in both the cell membrane and nucleus (Fig. 2A). To investigate the expression levels of *FveDAD2* in different organs of the woodland strawberry, we performed Quantitative RT-PCR (qRT-PCR) analysis. The results showed that *FveDAD2* was expressed in all tested organs. The highest transcript abundance was detected in the young leaf, followed by the shoot tip, flower, fruit, mature leaf, root, and petiole (Fig. 2B).

**Figure 2.**
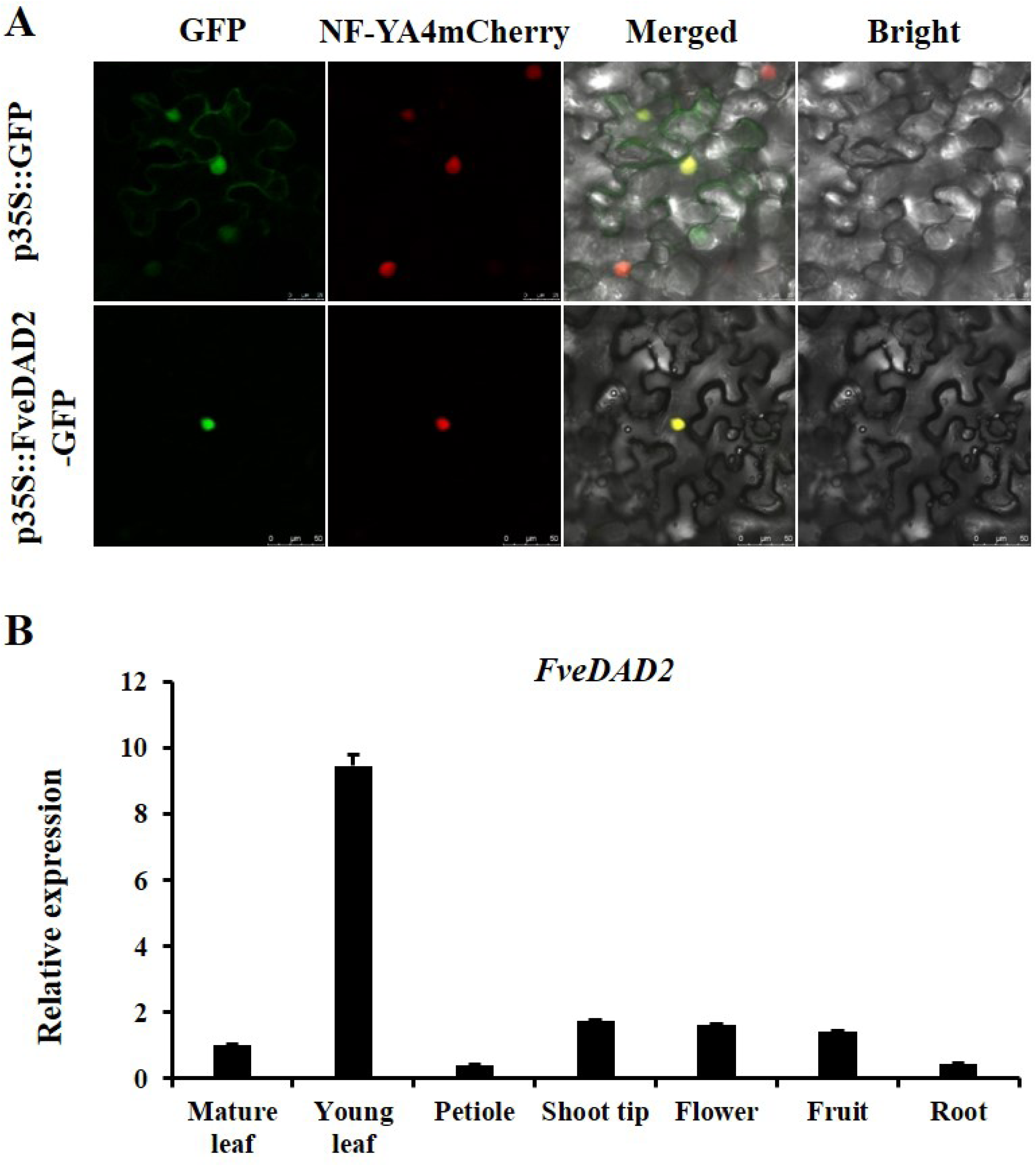
Subcellular localization and expression pattern of *FveDAD2* **A)** Subcellular localization was observed for a *pro35S::FveDAD2-GFP* fusion protein in *Nicotiana benthamiana* leaves. Bar = 50 μm. **B)** qRT-PCR analysis was conducted for the transcript levels of *FveDAD2* in various organs during the reproductive growth phase.

### Silencing of *FveDAD2* increased the number of branch crowns in Yellow Wonder

To elucidate the role of *FveDAD2* in the woodland strawberry, we utilized *Agrobacterium*-mediated transformation to introduce FveDAD2-RNAi vectors into the woodland strawberry accession ‘Yellow Wonder’ (YW). Three FveDAD2-RNAi lines (#1, #2, #3) were obtained (Fig. 3, A and B). We examined these RNAi silencing lines at the DNA level (Fig. 3A) and the RNA level. The expression of *FveDAD2* in RNAi lines was decreased to one-tenth of the YW (Fig. 3B). The phenotypes of RNAi lines and the YW were recorded during vegetative stages (Fig. 3C). All FveDAD2-RNAi lines had more branch crowns than YW. We also identified some phenotypic differences between FveDAD2-RNAi lines and YW, and the results showed that plants of FveDAD2-RNAi lines exhibited shorter plant height, increased number of leaves, and reduced area of the third leaf (Fig. 3, D-G). Thus, FveDAD2 plays a negative role in regulating branch crowns. In addition, we found that the FveDAD2-RNAi lines had significantly smaller fruits than YW (Fig. 3H), with a 26% reduction in single fruit weight (Fig. 3I) and an 11% reduction in total soluble solids content (Fig. 3J) in the FveDAD2-RNAi fruits compared to the fruits of YW.

**Figure 3.**
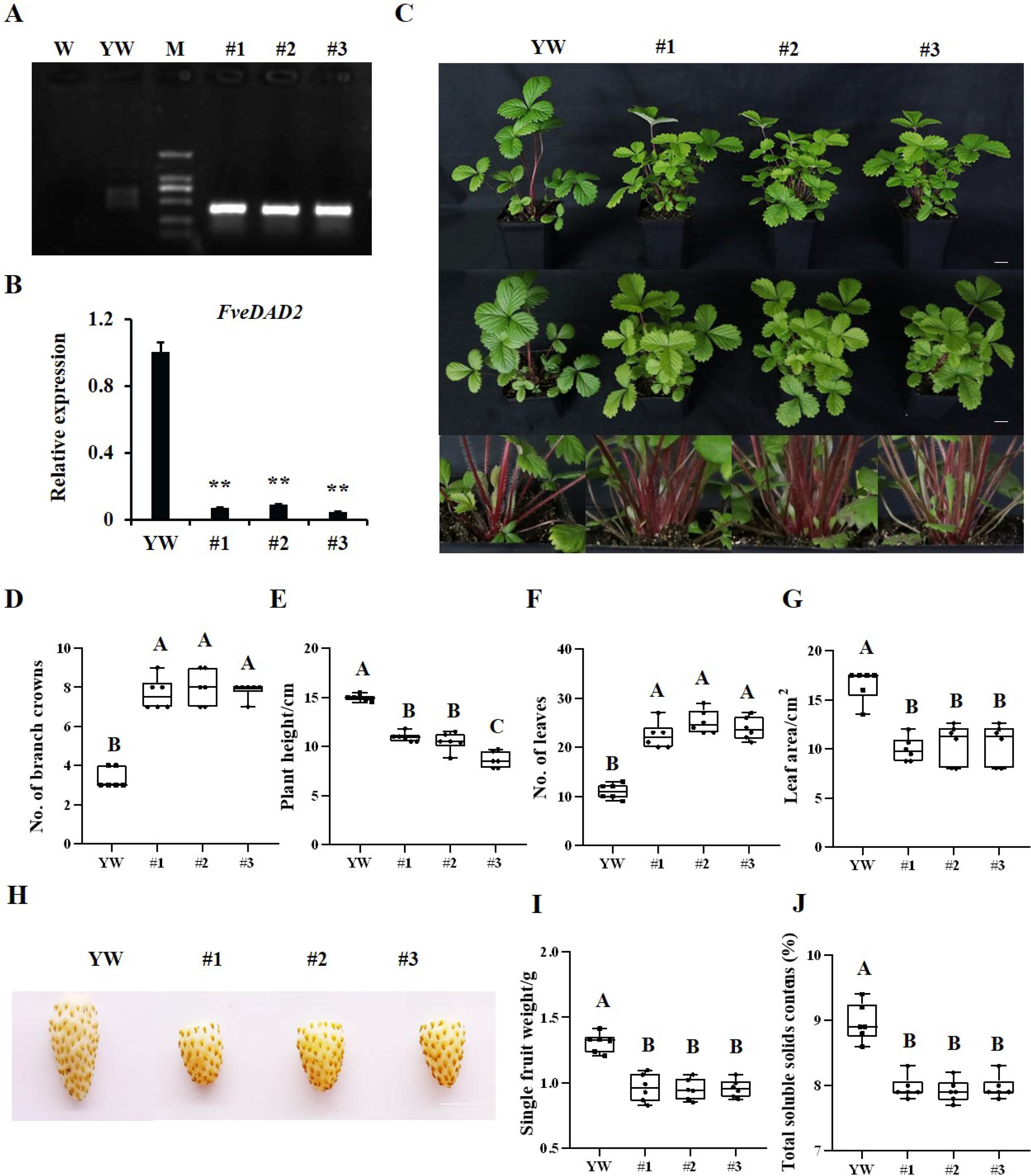
The phenotype of *FveDAD*2 silencing in YW **A)** PCR analysis of the *FveDAD2* in YW and FveDAD2-RNAi lines. W: water; YW: non-transgenic control; M: DL2000 marker; #1, #2, #3: three independent positive FveDAD2-RNAi transgenic lines. **B)** Analysis of relative expression of *FveDAD2* by qRT-PCR in YW and FveDAD2-RNAi lines (DPS, Duncan’s MRT, \*\**P* < 0.01). **C)** The phenotypes of FveDAD2-RNAi lines and the YW during the vegetative stage. Bar = 2 cm. The phenotypes of fruit **H)** in the FveDAD2-RNAi lines and the YW. Bar = 1 cm. The quantification of the number of branch crowns **D)**, plant height **E)**, the number of leaves **F)**, the area of the third leaf **G)**, the single fruit weight **I)**, and the total soluble solids contents **J)**. Values are Mean ± SD (*n* = 6), A–C indicates significant differences among the plants (Prism 8.0.1, Duncan’s MRT, *P* < 0.01).

### *FveDAD2* regulated branching-crowns-related genes mainly in hormones, macroelements, and sugars

To find the reason for *FveDAD2* regulating branch crowns in the strawberry, we performed transcriptome analysis of shoot tips in FveDAD2-RNAi and YW. There were 204 different expression genes (DEGs; Fold Change ≥ 1.5 and FDR < 0.05). Among them, 106 were upregulated and 98 were downregulated in the FveDAD2-RNAi plants compared with YW (Fig. 4A). DEGs in RNAi plants and YW enriched in plant hormone signaling, starch and sucrose metabolism according to KEGG analysis (Fig. 4B). Next, we conducted the NR annotations of DEGs and screened the genes related to branching, and found that these genes accounted for about 25% of the total DEGs. These branching-related genes were mainly involved in hormones, macroelements, and sugars. (Fig. 4C).

**Figure 4.**
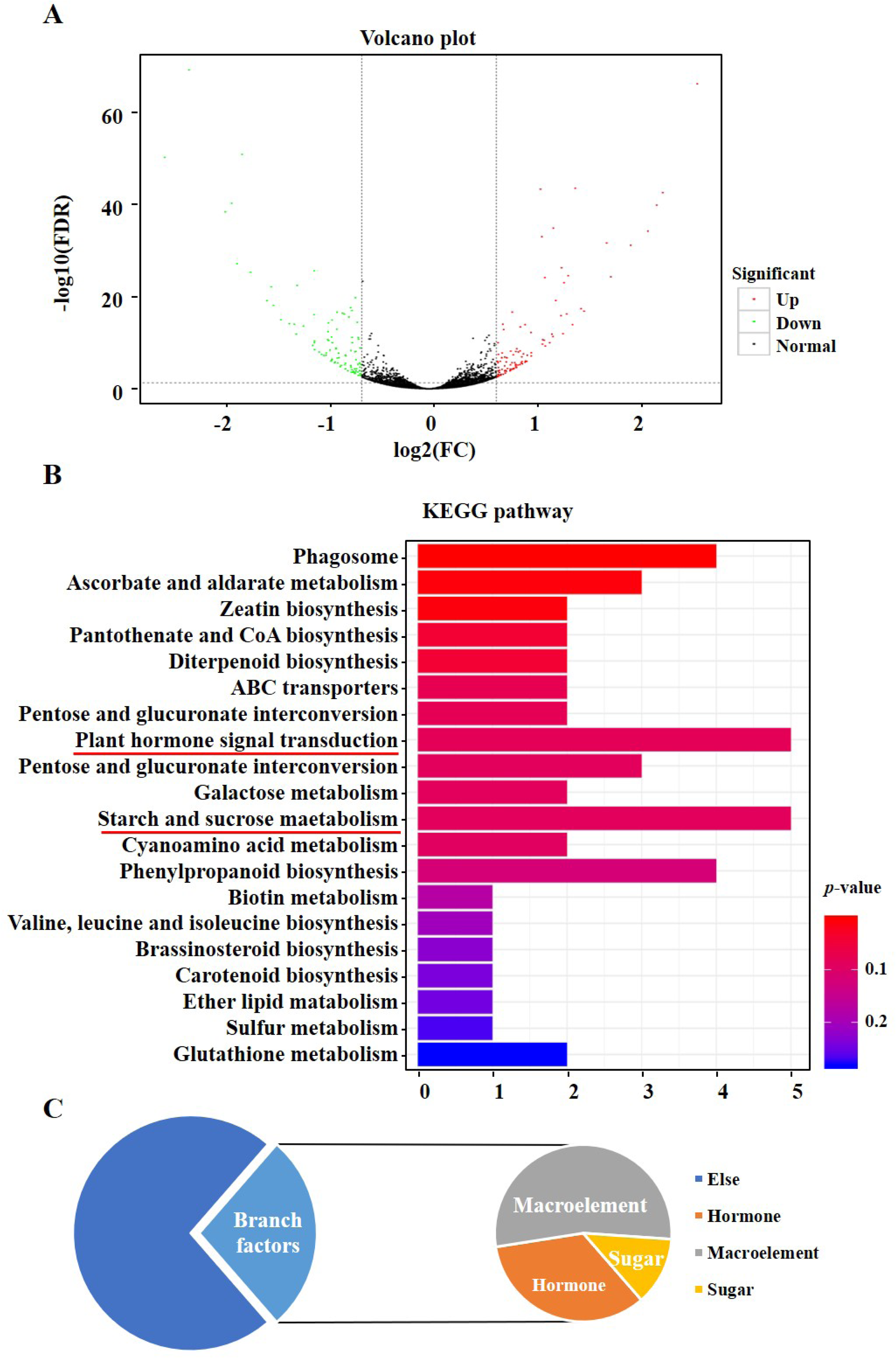
RNA-seq analysis of shoot tips in *FveDAD2*-RNAi transgenic lines and YW **A)** Differential expression volcano plot in the transcriptome data (Fold Change ≥ 1.5 and FDR < 0.05). The green points are the down-regulated DEGs, the red is the up-regulated DEGs, and the black is the non-DEGs. **B)** KEGG enrichment of genes in the transcriptome analysis. The red line indicates the pathway where DEGs are enriched to a greater extent. **C)** Identifying the DEGs related to branching through the NR annotation.

### FveDAD2 interacts with FveSMXL7 depending on the SL

SL binds to the D14/DAD2 receptor, which recruits the D3/MAX2 and D53/SMXL6,7,8 to form an SCF complex. As a result, D53/SMXL6,7,8 undertakes proteasomal degradation, leading to the SL signaling response (Waters et al., 2017). We hypothesize that FveDAD2 interacts with the downstream repressor to transduce SL signals. Then, the *FveSMXL7* was identified by searching for homologous genes in the woodland strawberry in the GDR, using the rice *OsD53* mRNA sequence. This gene was found to have a high similarity of 50.2% to *AtSMXL7* in the Tair database.

We cloned the CDS of *FveSMXL7* and performed amino acid comparisons with OsD53 and AtSMXL6/7/8. FveSMXL7 is recognized by four domains: N-terminal domain, D1 ATPase domain, M domain, and D2 ATPase domain (Fig. 5A; Supplemental Fig. S2). For D14 interaction, D1 is a necessary and sufficient condition. The D2 is the key domain of the receptor-induced degradation (Khosla et al., 2020). Notably, there are four protein domains, two Walker A (WA) and two Walker B (WB) in the D1 and the D2 domains, which are important parts of the structural domain of the ClpB protein superfamily (Zhou et al., 2013; Kerr et al., 2021). Additionally, the D2 domain of FveSMXL7 contains a highly conserved RGKT motif, found in D53 and AtSMXL6/7/8 and thought to be vital for SL-mediated D53/SMXLs protein degradation; and a highly conserved sequence, LDLNL, which is the Ethylene responsive element binding factor-associated Amphiphilic Repression (EAR) motif, which is known to determine whether a TF is a repressor (Kagale and Rozwadowski, 2011; Jiang et al., 2013; Zhou et al., 2013; Soundappan et al., 2015; Wang et al., 2015). The comparisons revealed that FveSMXL7 is structurally conserved and consistent with the model plant.

**Figure 5.**
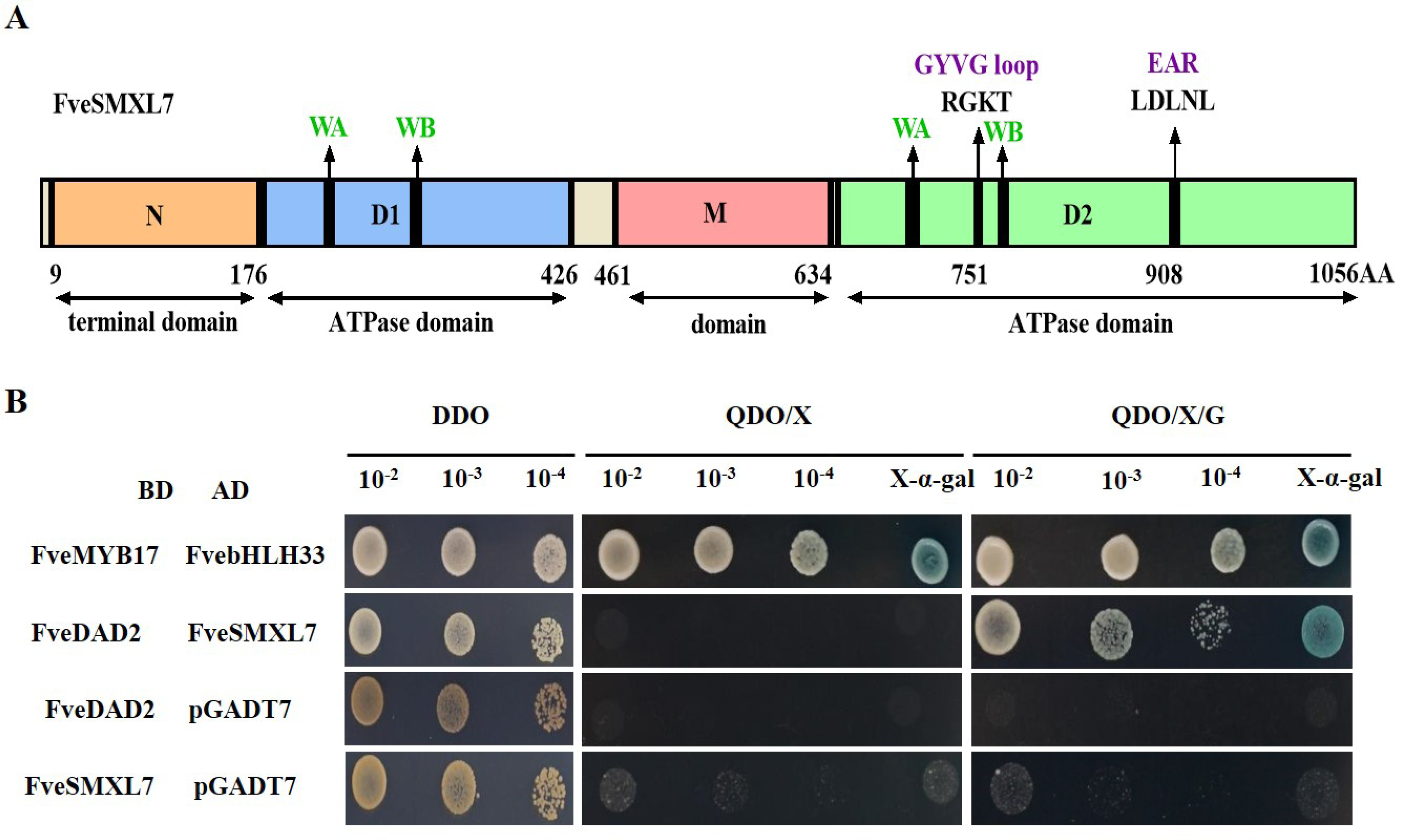
Interaction of FveDAD2 with FveSMXL7 depends on GR24 **A)** The structure of the FveSMXL7 protein displays four protein domains and two essential amino acid motifs: two Walker A (WA), two Walker B (WB), GYVG loop, and EAR motifs. **B)** The Y2H assay demonstrated that FveDAD2 can interact with FveSMXL7 in the presence of GR24. The media of this experiment involves the minimal media double dropouts (DDO, SD-Leu/-Trp), the minimal media quadruple dropouts (QDO/X, SD-Leu/-Trp/-His/-Ade plus X-a-gal), and (QDO/X/G, SD-Leu/-Trp/-His/-Ade plus X-a-gal and GR24). Binding domain (BD), activating domain (AD), and synthetic dropout (SD) were used.

In the presence of GR24, D53 may interact with D14 (Zhou et al., 2013). We investigated the interaction between FveDAD2 and FveSMXL7 using yeast two-hybrid (Y2H) and observed physical interactions between FveDAD2 and FveSMXL7. This interaction was dependent on GR24 (Fig. 5B). These results suggest that FveDAD2 and FveSMXL7, as members of the SL signaling pathway, are structurally and functionally conserved, which provides a solid foundation to explore downstream components of SL-regulated strawberry branch crowns.

### FveSMXL7 indirectly promotes *FveABA8’OH1* expression through the transcription factor FveHB7

To further reveal the molecular mechanism of the SL signaling pathway regulating branch crowns of the woodland strawberry, two additional components related to ABA were identified based on the transcriptome data of FveDAD2-RNAi and YW. One down-regulated gene encodes a TF, the homeodomain-leucine zipper (HD-Zip) class I HB7, which has a crucial function in the ABA signaling pathway (Ré et al., 2010; Son et al., 2010; Valdes et al., 2012; Zhao et al., 2021).

It was found that SMXL6 and SMXL7 can directly bind to promoters with ATAACAA motif and negatively regulate gene expression (Wang et al., 2020). We analyzed the promoter of *FveHB7* and found a possible binding motif ATAACAA at 1,642 bp upstream of the transcription start site. The Y1H assay suggested that FveSMXL7 could bind to the promoter of *FveHB7* (Fig. 6A). We next examined the effect of FveSMXL7 on the transcription of *FveHB7* utilizing the luciferase reporter system and the β-glucuronidase (GUS) transactivation assay. When *pro35S::FveSMXL7* was co-transformed with the *proFveHB7::LUC*, the LUC signal was significantly weakened or even disappeared compared with the control (Fig. 6B). When *pro35S::FveSMXL7* was co-transformed with the *proFveHB7::GUS*, the GUS activity demonstrated significant reduction in comparison to the control (Fig. 6D). These data show that FveSMXL7 repressed the expression of the *FveHB7*.

**Figure 6.**
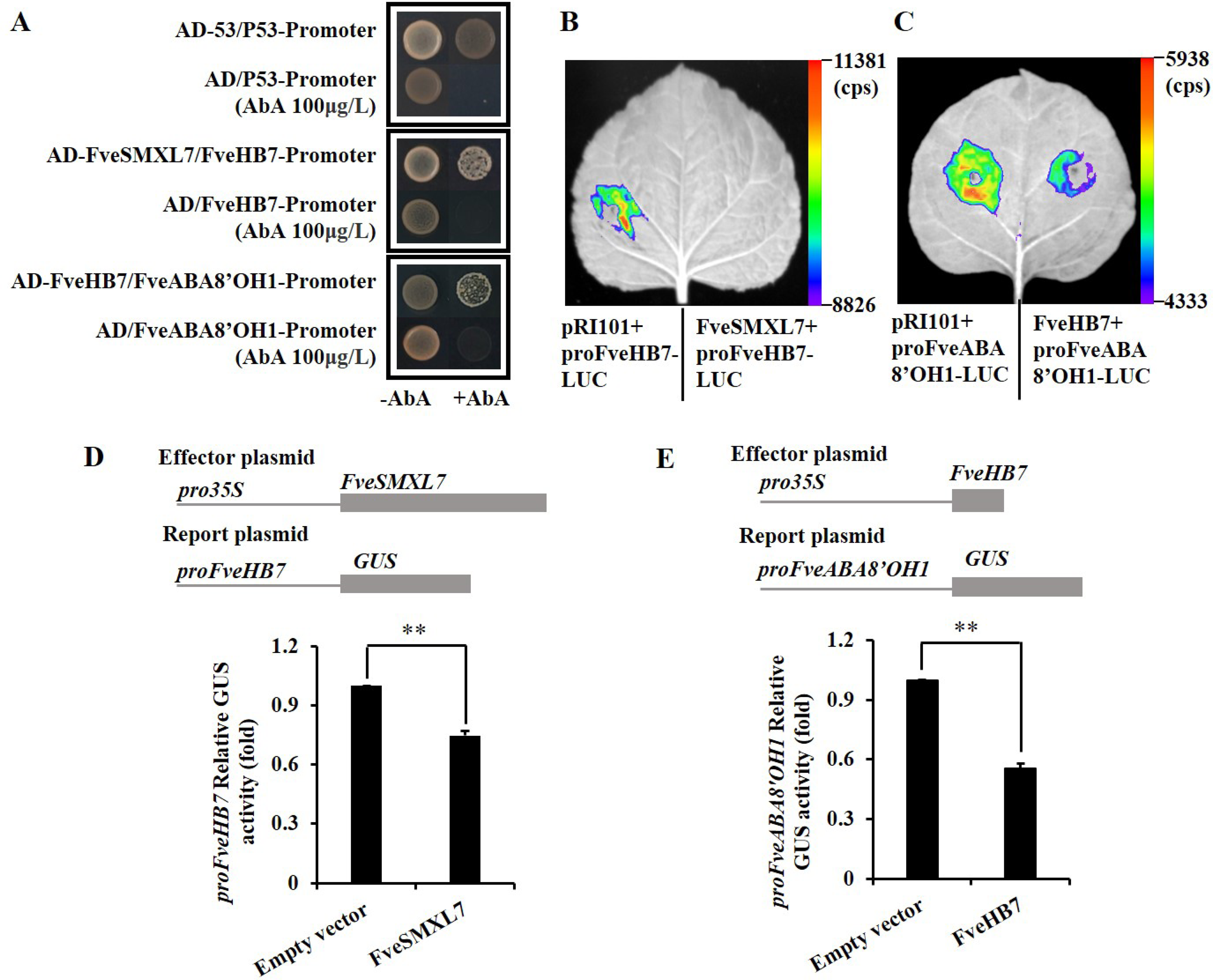
FveSMXL7 indirectly promotes *FveABA8’OH1* expression via the transcription factor FveHB7. **A)** Y1H assay indicates that FveSMXL7/FveHB7 binds to the *FveHB7*/*FveABA8’OH1* promoter. The baseline concentration of aureobasidin A (AbA) was 100 μg/L. The positive controls utilized were AD-53 and the P53 promoter. The negative controls utilized were the empty vector and the *FveHB7*/*FveABA8’OH1* promoter. **B)** and **C)** Luciferase reporter assay. *pro35S::FveSMXL7/FveHB7* and *proFveHB7/FveABA8’OH1::LUC* were cotransformed into *Nicotiana benthamiana*. **D)** and **E)** GUS activity analysis. *pro35S::FveSMXL7/FveHB7* and *proFveHB7/FveABA8’OH1::GUS* were cotransformed into *N. benthamiana* (DPS, Duncan’s MRT, \*\**P* < 0.01).

Three HD-Zip family proteins and BRC1 promote the expression of *NCED3*, which participates in ABA biosynthesis. This leads to an increase in ABA content, which inhibits bud formation (González-Grandío et al., 2016). However, the expression of *NCEDs* was not significantly different in the transcriptome data of the FveDAD2-RNAi line. An up-regulated key ABA catabolic enzyme *Abscisic acid 8’-hydroxylase 1* (*ABA8’OH1/ CYP707A*) was identified, which modulates endogenous ABA homeostasis in the many plants (Krochko et al., 1998; Li et al., 2015; Kim et al., 2020; Wang et al., 2023). We hypothesize that FveHB7 affects ABA levels through FveABA8’OH1. In tomatoes, the HD-zip family TF, SlHB15A, has been discovered to bind to an ATGAT DNA motif, which alters the cross-talk between auxin, jasmonic acid (JA), and ethylene that regulates flower pedicel abscission and fruit set (Clepet et al., 2021; Liu et al., 2022). As expected, we found three possible binding sites (ATGAT) at -595 bp, -205 bp, and -82 bp in the promoter of *FveABA8’OH1*. We cloned the promoter of *FveABA8’OH1* with a length of 742 bp and conducted Y1H assays. The Y1H results showed that FveHB7 can bind to the promoter of *FveABA8’OH1* (Fig. 6A). LUC and GUS assays were carried out to investigate further whether FveHB7 negatively regulates the transcription of *FveABA8’OH1* (Fig. 6, C and E). The results show that FveHB7 can bind to the promoter of *FveABA8’OH1* and repress the expression of *FveABA8’OH1*.

The above results indicate that FveSMXL7 binds to the *FveHB7* promoter and negatively regulates its expression, whereas FveHB7 binds to the *FveABA8’OH1* promoter and represses its transcriptional activity. In other words, FveSMXL7 indirectly enhances *FveABA8’OH1* expression by suppressing the transcript abundance of *FveHB7*.

### ABA levels were reduced in the shoot tips of the FveDAD2-RNAi lines

To verify whether ABA is associated with the regulation of strawberry branch crowns. The ABA levels in the shoot tips of both FveDAD2-RNAi lines and non-transgenic plants were analyzed using 60-day growing plants under comparable growth conditions. The FveDAD2-RNAi lines exhibited 2-3 branch crowns (red arrow), whereas the YW lines did not possess branch crowns. The transgenic lines exhibited a distinct branching and dwarfing phenotype in contrast to the YW (Fig. 7A). We found the content of ABA in *FveDAD2*-RNAi decreased by approximately 40% compared with that in YW (Fig. 7B). The findings above provide evidence that the reduction in ABA levels leads to a multi-branched phenotype in FveDAD2-RNAi lines.

**Figure 7.**
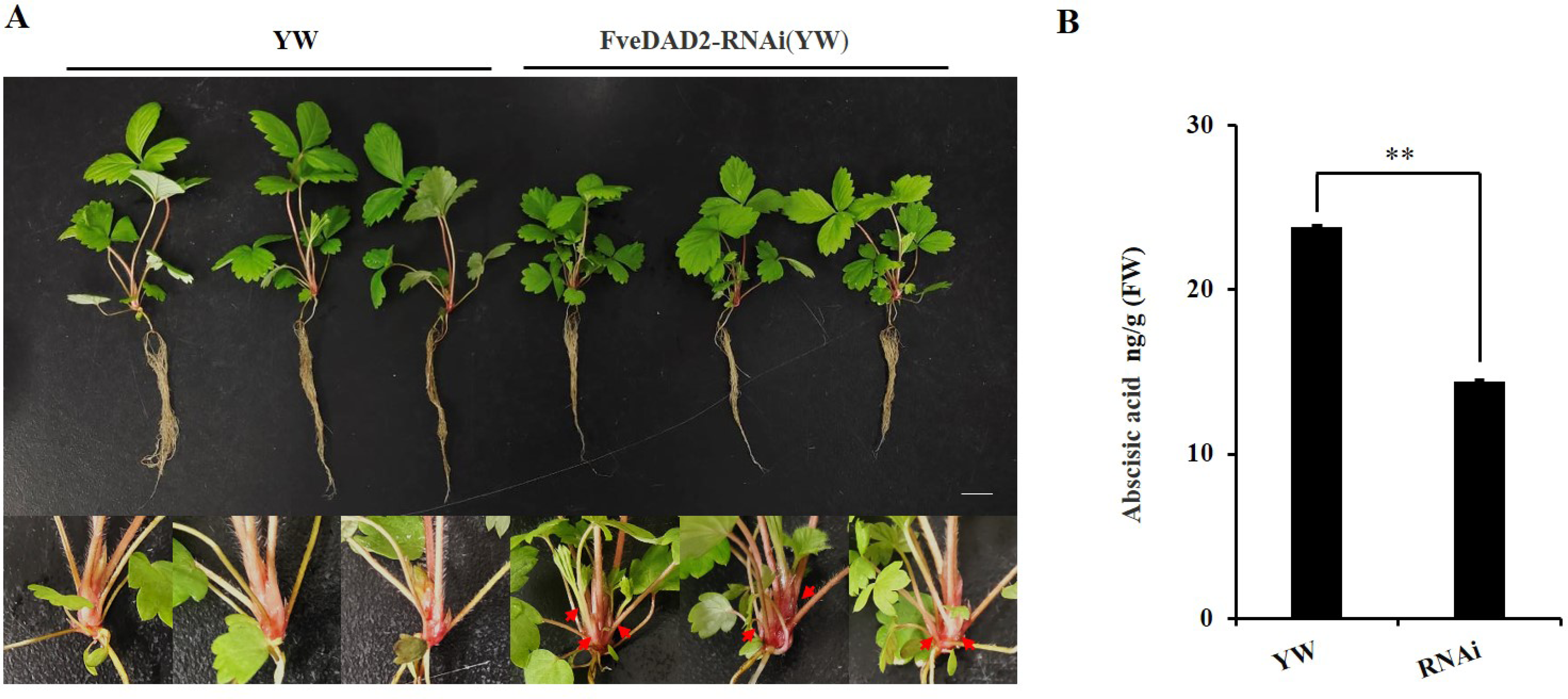
Abscisic acid was detected in the shoot tips of FveDAD2-RNAi lines and YW. **A)** Plant morphology in phytohormone assays using live seedlings grown for 60 days under comparable conditions. Bar = 1cm. **B)** There are absentic acid (ABA) levels in the shoot tips of FveDAD2-RNAi lines and YW.

To further determine the regulatory effect of ABA on strawberry branch crowns, we treated woodland strawberry YW with ABA and fluridone (an inhibitor of ABA) for a month. The ABA-treated YW plants significantly suppressed the number of crown branches, whereas YW plants with fluridone treatment showed an increased number of crown branches (Fig. 8, A and B). Significant leaf number and area differences were also observed among different treatments (Fig. 8, C and D). These data further support the idea that ABA content in shoot tips is a negative regulator of the strawberry crown branch.

**Figure 8.**
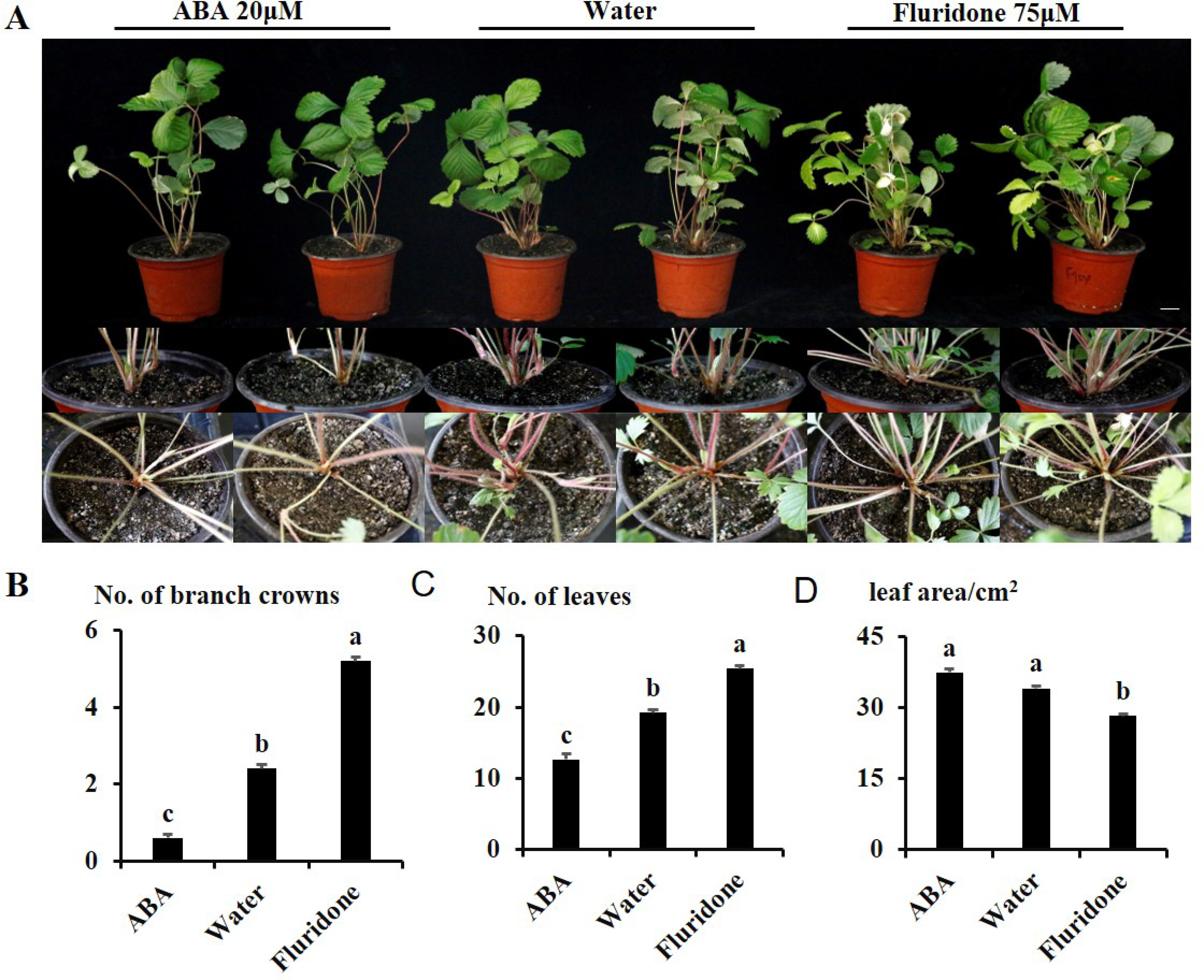
Effect of ABA and ABA inhibitor on branch crowns and leaves in YW. **A)** Phenotypes of YW after one month of treatment with ABA and fluridone (an inhibitor of ABA), Bar = 2cm. Number of branch crowns **B)**, number of leaves **C)**, and leaf area **D)** after one month of ABA and fluridone treatment in YW. Values are Mean ± SD (*n* = 5), a–c indicate significant differences among the plants (DPS, Duncan’s MRT, *P* < 0.05).

## Discussion

### SL receptors in woodland strawberry

We placed the SL receptor in a branch crowns model of the woodland strawberry. We searched the GDR database using the amino acid sequence of AtD14 to retrieve three homologous α/β hydrolases, which were FvH4_2g33160.t1, FvH4_1g17110.t1, and FvH4_1g05320.t1, respectively, based on the smallest to the largest *E*-value and corresponding to the FveDAD2 (XP_004290965.1), FveKAI2 (XM_004287925.2), and FveD14 (XM_004287028.2) in the NCBI database. Arabidopsis has three members of the alpha/beta hydrolase protein family. DWARF14 (D14) plays an essential role in SL signaling. The D14 homolog, KARRIKIN-INSENSITIVE 2 (KAI2), facilitates signaling through exogenous karrikins (KARs) or unknown endogenous KAI2 ligands (KL). Additionally, both KARs and SL have butenolide chemical structures, and genetic research indicates that MAX2 serves as a common protein for KAI2- and D14-mediated signaling pathways (Li and Tran, 2015; Mostofa et al., 2018). In addition, SMAX1 and SMXL2, two homologs of SMXL6/7/8, negatively regulate the KAI2-mediated signaling (Stanga et al., 2013; Stanga et al., 2016; Li et al., 2020). There is a third family member, DWARF 14-LIKE2 (DLK2), in *Arabidopsis*, which is structurally similar to KAI2 and D14. Unlike KAI2 and D14, DLK2 is not subject to rac-GR24-induced degradation and does not affect SL responses. DLK2 may function independently of MAX2 (Waters et al., 2012; Hu et al., 2021).

Alignment of the amino acid sequence of three homologous proteins with AtD14 (Supplemental Fig. S1) revealed that all three proteins in strawberries possess catalytic triads (Ser97-S^AtD14^, Asp218-D^AtD14^, His247-H^AtD14^) with hydrolase characteristics (Waters et al., 2012; de Saint Germain et al., 2016; Yao et al., 2016; Vegh et al., 2017; Yao et al., 2018). FveD14 lacks the conserved amino acid residues Gly159-G^AtD14^, Pro162-P^AtD14^, Glu175-E^AtD14^, and Ser177-R^AtD14^ in contrast to FveDAD2 and FveKAI2. The interaction between AtD14 and D3 requires Gly159-G^AtD14^, Pro162-P^AtD14^, and Glu175-E^AtD14^ in *Arabidopsis* (Vegh et al., 2017). Notably, Ser177-R^AtD14^ plays a crucial role in the binding of SsD14 to SMXL7 in sugarcane (Hu et al., 2021).

We further used the yeast two-hybrid assay to differentiate the three homologous proteins in the strawberry. We found that FveD14 lacking Ser177-R^AtD14^ failed to interact with SMXL7 with or without the addition of GR24, whereas both FveDAD2 and FveKAI2 interacted with SMXL7 in the presence of GR24, consistent with previous findings (Fig. 5B; Supplemental Fig. S3A). Based on the above experimental evidence, we tentatively assigned FveDAD2, FveD14, and FveKAI2 corresponding to AtD14, AtDLK2, and AtKAI2, respectively.

### FveSMXL7 indirectly enhances the expression of FveABA8’OH1 via the transcription factor FveHB7

D53/SMXL6,7,8 are transcriptional regulators that lack direct DNA binding ability, suggesting that they require adaptors to bind DNA to facilitate SL-regulated transcription and shoot branching (Hu et al., 2020). D53 interacts through the EAR motif with TPR2 to recruit TFs that regulate SL signaling and shoot branching. In this study, based on transcriptomic data of FveDAD2-RNAi and YW, a down-regulated transcription factor, FveHB7, associated with the ABA signaling pathway caught our attention. We examined the interactions between FveSMXL7 and FveHB7 using Y2H (Supplemental Fig. S3B) and found that they did not interact. Moreover, it was found that SMXL6 and SMXL7 can directly bind to the promoters of *SMXL6/7/8* and repress their transcription. The sequence ATAACAA is crucial for direct binding and repression (Wang et al., 2020). Interestingly, the *FveHB7* promoter contains an ATAACAA sequence, and our data show that FveSMXL7 binds to the *FveHB7* promoter and negatively regulates its expression (Fig. 6, A B and D).

Transcriptional targets of the SL pathway for shoot growth regulation, *TB1/FCL1/BRC1*, are linked to ABA. *BRC1* promotes the expression of three TFs (HB21, HB40, and HB53), which stimulate the expression of *NCED3*, resulting in ABA accumulation and subsequently affecting meristematic conditions (González-Grandío et al., 2016). Our data shows that the downstream regulation of FveDAD2 and FveSMXL7 in the strawberry may be independent of BRC1 or may not impact the expression of *NCED3*, a downstream target of BRC1. However, these components (*BRC1* and *NCED3*) were not found in the DEGs of our transcriptomic data between FveDAD2-RNAi and YW. Interestingly, we identified another ABA catabolic enzyme, FveABA8’OH1. The HD-Zip family TFs can bind to a specific DNA sequence, CAAT(A/T)ATTG, and promote gene expression (Shao et al., 2018; Zhang et al., 2021; Zhao et al., 2021; Dong et al., 2022). However, we did not find the CAAT(A/T)ATTG site for FveHB7 in the promoter fragment of *FveABA8’OH1*. It was found that the HD-Zip family can also bind to the ATGAT DNA motifs and repress expression of the target gene, and we found three such binding motifs, ATGAT, in the promoter of *FveABA8’OH*. Indeed, Y1H, Luciferase reporter assay, and GUS analysis showed that FveHB7 negatively regulates the promoter activity of *FveABA8’OH1* (Fig. 6, A C and E). Taken together, we identified a novel pathway for SL signaling.

### *FveDAD2* and *FveSMXL7* can be candidate target genes for genetic improvement of strawberry plant and fruit size

It is well known that plant branching is inextricably linked to yield, and the multi-branched phenotype of FveDAD2-RNAi transgenic plants was manifested not only during the vegetative period but also during the reproductive period (Supplemental Fig. S5A). In contrast to the wild type, the transgenic flowers were significantly smaller (Supplemental Fig. S5, B and C), produced smaller fruits (Fig. 3, H and I), and had a significantly reduced soluble solids content (Fig. 3J). Studies on the expression of SL-related genes in strawberry fruits at different developmental phases may help to explain this phenotype. The expression of SL pathway genes and SL concentration were higher in early fruit development but decreased as fruit development and ripening progressed, suggesting that SL is involved in the early development of strawberry fruits (Wu et al., 2019), which is in line with our results. The expression of both *FveD27* and *FveSMXL7* was reduced in the FveDAD2-RNAi transgenic plants (Supplemental Fig. S4, A, and B), and it was hypothesized that SL might affect the development of strawberry fruits during the green fruit expansion period. In addition, the FveABA8’OH1 homolog, FveCYP707A4a, has been linked to regulating the fruit size and ripeness through the control ABA levels and coordinating auxin, GA, and ABA signaling (Liao et al., 2018). The reduced fruit size phenotypes of the FveDAD2-RNAi plants indicate that SL may participate in the complex network of multiple hormones that regulate plant branching and fruit morphology. *FveDAD2* and *FveSMXL7* may be candidate target genes for the genetic improvement of strawberry plants and fruit size.

## Conclusion

We propose a model for regulating the branch crowns by FveDAD2 in the woodland strawberry. In the wild-type plants, FveDAD2 interacts with FveSMXL7 and ubiquitinates FveSMXL7 (Fig. 9A), whereas downregulation of the *FveDAD2* results in reduced ubiquitination of FveSMXL7, leading to increased repression of *FveHB7* transcription by FveSMXL7. This repression also results in reduced negative regulation of the *FveABA8OH1* by FveHB7. The increased *FveABA8OH1* transcript level decreased ABA accumulation in the shoot tips, ultimately increasing crown branching (Fig. 9B).

**Figure 9.**
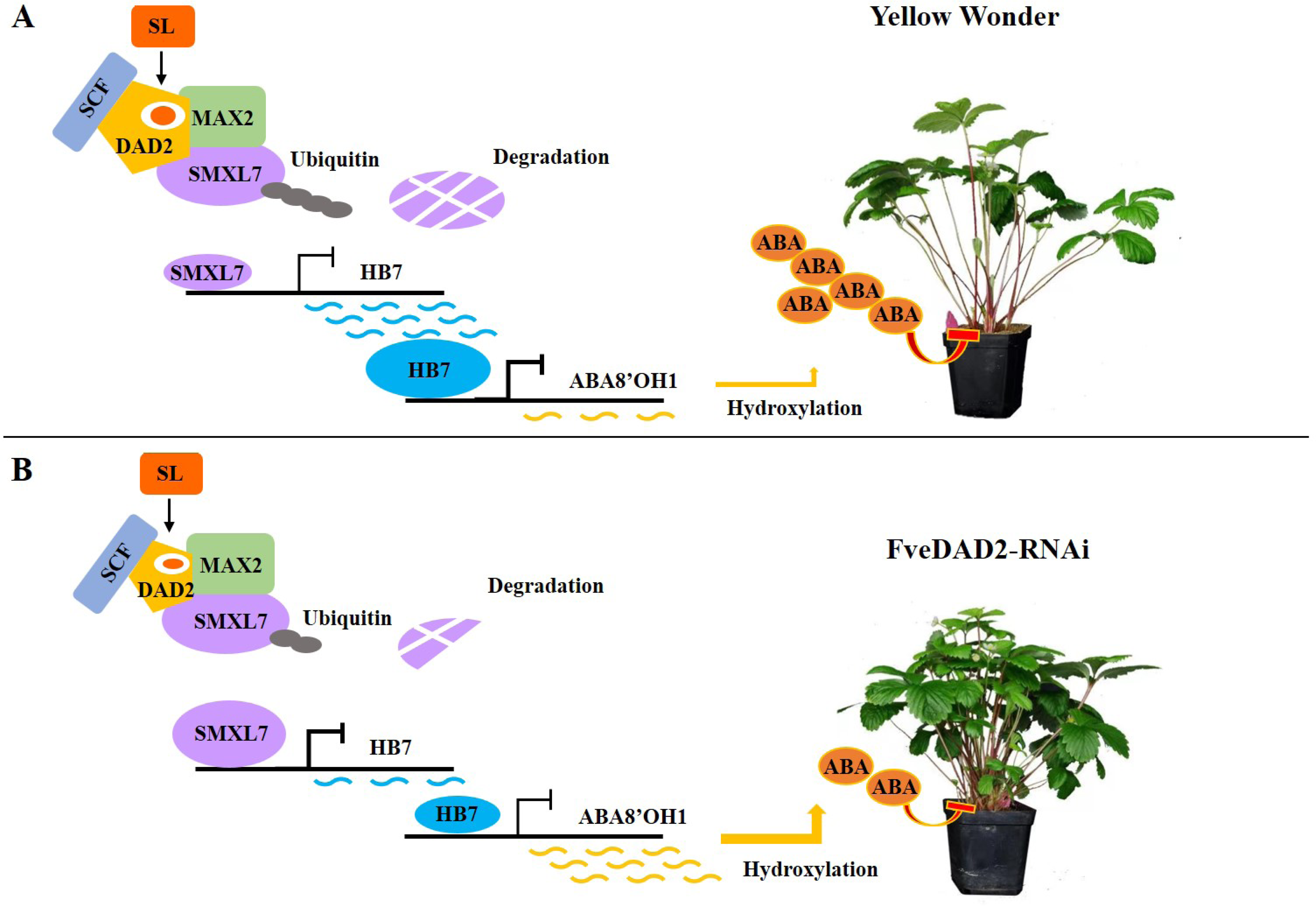
The model of SL signaling to regulate branch crowns by affecting ABA Metabolism in the woodland strawberry. FveDAD2 negatively regulates branch crowns by affecting abscisic acid levels by interacting with FveSMXL7 to regulate the expression of FveABA8’OH1 via the transcription factor FveHB7. **A)** YW, **B)** FveDAD2-RNAi line.

## Materials and Methods

### Phylogeny and sequence analysis

The phylogenetic tree of DAD2 proteins from various plants was constructed using MEGA6.0. The neighbor-joining (NJ) method was used to construct the tree with bootstrap analysis of 1000 replications. The FveDAD2 orthologs used for the tree construction were obtained from 44 species of higher plants. (Table S1). Conservative domain analysis of FveDAD2 and homologous proteins from 10 species of monocots (*Hordeum vulgare*, *Oryza sativa*, *Sorghum bicolor*, *Triticum aestivum*, *and Zea mays*) and dicots (*Arabidopsis thaliana*, *Fragaria vesca*, *Glycine max*, *Petunia hybrida*, *and Pisum sativum*) was conducted by the MEME online website (https://meme-suite.org/meme/doc/meme.html). DANMAN5.0 conducted protein sequence alignment.

### Plant materials and treatments

The diploid woodland strawberry Yellow Wonder (YW) was cultured under greenhouse conditions (16 h/8 h, light/dark, 23 °C) at Shenyang Agricultural University, China, using substrates (an appropriate ratio of grass charcoal, vermiculite, earthworm manure, and perlite). Different organs were harvested simultaneously for gene expression analysis during the early fruit ripening, frozen in liquid nitrogen, and stored at –80 °C.

*FveDAD2*-RNAi transgenic plants were obtained by the *Agrobacterium*-mediated leaf disc transformation(Li et al., 2018), which was performed using ‘YW’ at the tissue culture phase.

The *N. benthamiana* grown under a long-day photoperiod (16h/8h, light/dark) at 23°C for 30-50 days was used for transient expression analysis.

The ABA and fluridone-an ABA inhibitor (Coolaber, Beijing, China) treatments selected 50-day-old live YW seedlings grown in substrate that had not yet begun to branch and required consistent growth conditions and plant morphology. The experimental treatments contained ABA at 5, 10, and 20 μmol/L concentrations and fluridone at 25, 50, and 75 μmol/L concentrations. Water was the control group, with five plants in each group. The shoot tips were given 200 μL/time, administered twice weekly for one month.

### Gene expression analyses

The quantitative RT-PCR (qRT-PCR) was performed with SYBRGreen II (Takara, Dalian, China). The 7500 system (Applied Biosystems, Foster City, USA) was used for the analysis. The experiments were performed in three biological replicates, and the results were normalized using the housekeeping gene (Li et al., 2018). Different organs of YW and tender leaves from FveDAD2-RNAi plants were harvested for gene expression analysis. The transcription levels were estimated by the 2^−ΔΔCT^ method (Livak and Schmittgen, 2001). The primers involved are shown in Table S2.

### Subcellular localization

The CDS without the stop codon of *FveDAD2* was inserted into the pRI101-GFP to generate *pro35S::FveDAD2-GFP* by *KpnI* and *BamHI*. The *A. tumefaciens* strain GV3101 mediated the co-infiltration of corresponding vectors into the leaves of *N. benthamiana* (40 d old). Confocal fluorescence microscopy (Leica DMi8 A, Wetzlar, Germany) was used to observe the GFP fluorescence signal 48 h after transient transformation in the dark.

### Transcriptome sequencing analysis

The shoot tips of FveDAD2-RNAi and YW plants were harvested when there was a noticeable increase in branching before the reproductive growth period for transcriptome sequencing. The experiment involved six samples, each consisting of three individual plants, and three biological replicates were performed. The samples were sent to the Illumina sequencing platform at Biomarker Technologies (Beijing, China). Differentially transcribed genes were screened using a threshold of Fold Change ≥ 1.5 and FDR < 0.05. The genes were then functionally classified and analyzed differentially.

### Y2H assay

The Y2H assay was used to detect that FveDAD2 interacts with FveSMXL7. The CDS of *FveDAD2* was ligated into the pGBKT7 (BD) vector, and the CDS of *FveSMXL7* was inserted into the pGADT7 (AD) vector using the *EcoRI* and *BamHI*. Related materials and methods refer to previous methods(Gietz and Schiestl, 2007) and Saccharomyces cerevisiae competence preparation and transformation kit (Wuhan Protein Interaction Biotechnology Co., Ltd., Wuhan, China). The primers involved are shown in Table S2.

### Y1H assay

The Y1H assay confirmed that FveSMXL7 is bound to the promoter of FveHB7, and FveHB7 is bound to the promoter of *FveABA8’OH1*. The CDS of *FveHB7* was cloned into the pGADT7 vector with the *EcoRI* and *BamHI*. The *FveHB7* (1715 bp) promoter fragment and the *FveABA8’OH1* (742 b) promoter fragment were inserted into the pAbAi vector with the *HindIII* and *SacI*. Related materials and methods refer to previous methods(Gietz and Schiestl, 2007) and Saccharomyces cerevisiae competence preparation and transformation kit (Wuhan Protein Interaction Biotechnology Co., Ltd., Wuhan, China). The primers involved are shown in Table S2.

### Luciferase reporter assay

To generate the effector vector *pro35S::FveSMXL7* and *pro35S::FveHB7*, the CDS of *FveSMXL7* was ligated into the pRI101 vector through the *NdeI* and *BamHI*, and the CDS of *FveHB7* was inserted into the pRI101 vector through the *SalI* and *BamHI*. The *FveHB7*/*FveABA8’OH1* promoter sequence (2000 bp), respectively, was cloned into the pGreenII0800-LUC vector (Hellens et al., 2005) using the *HindΙΙΙ* and *BamHΙ*to construct the reporter vector. The *A. tumefaciens* strain GV3101(pSoup) mediated the co-infiltration of corresponding vectors into the leaves of *N. benthamiana* (50d old). LUC activity was observed as previously researched (Lei et al., 2020). The primers involved are shown in Table S2.

### GUS analysis

The effector vectors *pro35S::FveSMXL7* and *pro35S::FveHB7* have been constructed (refer to luciferase reporter assay). *FveHB7/FveABA8’OH1* promoter sequences (2000bp) were cloned into pRI201-GUS using *PstI* and *XbaI* to generate the reporter vectors. The *A. tumefaciens* strain GV3101 mediated the co-infiltration of corresponding vectors into the leaves of *N. benthamiana* (50d old). Measure of GUS activity in previous studies (Li et al., 2016). The primers involved are shown in Table S2.

### Hormone quantification

The shoot tips of YW and *FveDAD2*-RNAi cultured in perlite fixed with 1/2 Hoagland’s solution for 60 d (when differences in the number of branches began to appear) were selected as samples, and five plants were mixed. Each sample, approximately 0.5 g, was homogenized into a powder using liquid nitrogen. ABA was extracted and quantified according to the technical support (Wuhan MetwareBiotechnology Co., Ltd., Wuhan, China). The ABA was extracted by the Ultra Performance Liquid Chromatography (ExionLC™ AD, America) and Tandem Mass Spectrometry (SCIEX QTRAP 6500+, America) (LC-MS/MS) system. ABA was quantified by the external standard technique and is presented as ng/g fresh weight (FW).

### Plant phenotypic characterization statistics

*FveDAD2* transgenic plants and YW with uniform growth conditions were selected, and the following data were investigated and counted. Number of branches: a single branch with a central leaf was counted as a branch; plant height (cm): the vertical height from the base of the strawberry stem to the highest point of the plant; number of leaves: the strawberry leaf was counted as a single leaf when all the ternary leaves of the strawberry were expanded; leaf area (cm^2^): the product of the length and width of central leaf of the third ternately compound leaf from the center of the strawberry outward was calculated; the single fruit weight (g): the weight of a primary fruit; the total soluble solids contents of a primer fruit. To quantify the plant phenotypic characterization, A–C indicates significant differences among the plants (Prism 8.0.1 Duncan’s MRT, *n*=6, *P* < 0.01).

### Accession numbers

The Arabidopsis Information Resource (TAIR) https://www.arabidopsis.org/: AtD14 (AT3G03990.1), AtSMXL7 (AT2G29970.1), AtHB7 (AT2G46680.1); Genome Database for Rosaceae (GDR) https://www.rosaceae.org/: FveDAD2 (FvYW_2g33160.t1), FveKAI2(FvH4_1g17110.t1), FveD14(FvH4_1g05320.t1) FveSMXL7 (FvH4_7g24540.t1), FveHB7 (FvH4_7g17320.t1), and FveABA8’OH1 (FvH4_2g39460.t1); National Center for Biotechnology Information (NCBI) https://www.ncbi.nlm.nih.gov/: FveDAD2 (XP_004290965.1), FveKAI2(XM_004287925.2), FveD14(XM_004287028.2) OsD53 (XP_015617462.1), FveSMXL7 (XM_011471618.1), and FveHB7 (XP_004306630.1). The data of the phylogenetic tree and the conservative domain analysis are shown in Table S1.

## Supplemental Data

**Supplemental Table S1.** Sequence information for phylogenetic analysis of DAD2

**Supplemental Table S2.** All primer information included in this study

**Supplemental Figure S1.** Amino acid sequence comparison of FveDAD2 homologous proteins

**Supplemental Figure S2.** SMXL7 amino acid sequence comparison

**Supplemental Figure S3.** FveSMXL7-related yeast two-hybrid assay

**Supplemental Figure S4.** SL-related gene expression in FveDAD2-RNAi line

**Supplemental Figure S5.** Small flowers in the FveDAD2-RNAi line

## Acknowledgments

pGreenII0800-LUC has been kindly provided by Yuepeng Han of Wuhan Botanical Garden, Chinese Academy of Sciences.

## Author contributions

H. S., J. Z., and Z. Z. conceived and designed the experiments; H. S. and W. J. performed the experiments; H. S. and Y. W. analyzed the data and wrote the manuscript; all the authors revised and approved the final version.

## References

Aguilar-Martínez JA, Poza-Carrión Cs, Cubas P (2007) Arabidopsis BRANCHED1 acts as an integrator of branching signals within axillary buds. Plant Cell 19: 458–472

Barbier FF, Cao D, Fichtner F, Weiste C, Perez-Garcia MD, Caradeuc M, Le Gourrierec J, Sakr S, Beveridge CA (2021) HEXOKINASE1 signalling promotes shoot branching and interacts with cytokinin and strigolactone pathways. New Phytol 231: 1088–1104

Bertheloot J, Barbier F, Boudon F, Perez-Garcia MD, Péron T, Citerne S, Dun E, Beveridge C, Godin C, Sakr S (2019) Sugar availability suppresses the auxin-induced strigolactone pathway to promote bud outgrowth. New Phytol 225: 866–879

Braun N, de Saint Germain A, Pillot J-P, Boutet-Mercey S, Dalmais M, Antoniadi I, Li X, Maia-Grondard A, Le Signor C, Bouteiller N, Luo D, Bendahmane A, Turnbull C, Rameau C (2012) The Pea TCP Transcription Factor PsBRC1 Acts Downstream of Strigolactones to Control Shoot Branching. Plant Physiol 158: 225–238

Cao D, Chabikwa T, Barbier F, Dun EA, Fichtner F, Dong L, Kerr SC, Beveridge CA (2023) Auxin-independent effects of apical dominance induce changes in phytohormones correlated with bud outgrowth. Plant Physiol 192: 1420–1434

Chesterfield RJ, Vickers CE, Beveridge CA (2020) Translation of Strigolactones from Plant Hormone to Agriculture: Achievements, Future Perspectives, and Challenges. Trends Plant Sci 25: 1087–1106

Clepet C, Devani RS, Boumlik R, Hao Y, Morin H, Marcel F, Verdenaud M, Mania B, Brisou G, Citerne S, Mouille G, Lepeltier J-C, Koussevitzky S, Boualem A, Bendahmane A (2021) The miR166–SlHB15A regulatory module controls ovule development and parthenocarpic fruit set under adverse temperatures in tomato. Mol Plant 14: 1185–1198

Crawford S, Shinohara N, Sieberer T, Williamson L, George G, Hepworth J, Müller D, Domagalska MA, Leyser O (2010) Strigolactones enhance competition between shoot branches by dampening auxin transport. Development 137: 2905–2913

de Saint Germain A, Clavé G, Badet-Denisot M-A, Pillot J-P, Cornu D, Le Caer J-P, Burger M, Pelissier F, Retailleau P, Turnbull C, Bonhomme S, Chory J, Rameau C, Boyer F-D (2016) An histidine covalent receptor and butenolide complex mediates strigolactone perception. Nat Chem Biol 12: 787–794

Dong S, Tarkowska D, Sedaghatmehr M, Welsch M, Gupta S, Mueller-Roeber B, Balazadeh S (2022) The HB40-JUB1 transcriptional regulatory network controls gibberellin homeostasis in Arabidopsis. Mol Plant 15: 322–339

Dong Z, Xiao Y, Govindarajulu R, Feil R, Siddoway ML, Nielsen T, Lunn JE, Hawkins J, Whipple C, Chuck G (2019) The regulatory landscape of a core maize domestication module controlling bud dormancy and growth repression. Nat Commun 10: 3810

Duan J, Yu H, Yuan K, Liao Z, Meng X, Jing Y, Liu G, Chu J, Li J (2019) Strigolactone promotes cytokinin degradation through transcriptional activation of CYTOKININ OXIDASE/DEHYDROGENASE 9 in rice. Proc Natl Acad Sci U S A 116: 14319–14324

Fichtner F, Barbier FF, Feil R, Watanabe M, Annunziata MG, Chabikwa TG, Höfgen R, Stitt M, Beveridge CA, Lunn JE (2017) Trehalose 6-phosphate is involved in triggering axillary bud outgrowth in garden pea (Pisum sativum L.). Plant J 92: 611–623

Gietz RD, Schiestl RH (2007) High-efficiency yeast transformation using the LiAc/SS carrier DNA/PEG method. Nat Protoc 2: 31–34

Gomez-Roldan V, Fermas S, Brewer PB, Puech-Pagès V, Dun EA, Pillot J-P, Letisse F, Matusova R, Danoun S, Portais J-C, Bouwmeester H, Bécard G, Beveridge CA, Rameau C, Rochange SF (2008) Strigolactone inhibition of shoot branching. Nature 455: 189–194

González-Grandío E, Pajoro A, Franco-Zorrilla JM, Tarancón C, Immink RGH, Cubas P (2016) Abscisic acid signaling is controlled by a BRANCHED1/HD-ZIP I cascade in Arabidopsis axillary buds. Proc Natl Acad Sci U S A 114: e245-e254

Hamiaux C, Drummond RS, Janssen BJ, Ledger SE, Cooney JM, Newcomb RD, Snowden KC (2012) DAD2 is an alpha/beta hydrolase likely to be involved in the perception of the plant branching hormone, strigolactone. Curr Biol 22: 2032–2036

Hellens RP, Allan AC, Friel EN, Bolitho K, Grafton K, Templeton MD, Karunairetnam S, Gleave AP, Laing WA (2005) Transient expression vectors for functional genomics, quantification of promoter activity and RNA silencing in plants. Plant Methods 1: 1–14

Hu A, Zhao Q, Chen L, Zhao J, Wang Y, Feng K, Wu L, Xie M, Zhou X, Xiao L, Ming Z, Zhang M, Yao R (2021) Identification of Conserved and Divergent Strigolactone Receptors in Sugarcane Reveals a Key Residue Crucial for Plant Branching Control. Front Plant Sci 12: 747160

Hu J, Ji Y, Hu X, Sun S, Wang X (2020) BES1 Functions as the Co-regulator of D53-like SMXLs to Inhibit BRC1 Expression in Strigolactone-Regulated Shoot Branching in Arabidopsis. Plant Commun 1

Jiang L, Liu X, Xiong G, Liu H, Chen F, Wang L, Meng X, Liu G, Yu H, Yuan Y, Yi W, Zhao L, Ma H, He Y, Wu Z, Melcher K, Qian Q, Xu HE, Wang Y, Li J (2013) DWARF 53 acts as a repressor of strigolactone signalling in rice. Nature 504: 401–405

Kagale S, Rozwadowski K (2011) EAR motif-mediated transcriptional repression in plants. Epigenetics 6: 141–146

Kelly JH, Tucker MR, Brewer PB (2023) The strigolactone pathway is a target for modifying crop shoot architecture and yield. Biology 12: 95

Kerr SC, Patil SB, de Saint Germain A, Pillot JP, Saffar J, Ligerot Y, Aubert G, Citerne S, Bellec Y, Dun EA, Beveridge CA, Rameau C (2021) Integration of the SMXL/D53 strigolactone signalling repressors in the model of shoot branching regulation in Pisum sativum. Plant J 107: 1756–1770

Khosla A, Morffy N, Li Q, Faure L, Chang SH, Yao J, Zheng J, Cai ML, Stanga J, Flematti GR, Waters MT, Nelson DC (2020) Structure–Function Analysis of SMAX1 Reveals Domains That Mediate Its Karrikin-Induced Proteolysis and Interaction with the Receptor KAI2. Plant Cell 32: 2639–2659

Kim HM, Park SH, Ma SH, Park SY, Yun C-H, Jang G, Joung YH (2020) Promoted ABA Hydroxylation by Capsicum annuum CYP707As Overexpression Suppresses Pollen Maturation in Nicotiana tabacum. Front Plant Sci 11: 583767

Krochko JE, Abrams GD, Loewen MK, Abrams SR, Cutler AJ (1998) (+)-Abscisic acid 8′-hydroxylase is a cytochrome P450 monooxygenase. Plant Physiol 118: 849–860.

Lei Y, Sun Y, Wang B, Yu S, Dai H, Li H, Zhang Z, Zhang J (2020) Woodland strawberry WRKY71 acts as a promoter of flowering via a transcriptional regulatory cascade. Hortic Res 7: 137

Li Q, Chen P, Dai S, Sun Y, Yuan B, Kai W, Pei Y, He S, Liang B, Zhang Y, Leng P (2015) PacCYP707A2 negatively regulates cherry fruit ripening while PacCYP707A1 mediates drought tolerance. J Exp Bot 66: 3765–3774

Li T, Jiang Z, Zhang L, Tan D, Wei Y, Yuan H, Li T, Wang A (2016) Apple (Malus domestica) MdERF2 negatively affects ethylene biosynthesis during fruit ripening by suppressingMdACS1transcription. Plant J 88: 735–748

Li W, Zhang J, Sun H, Wang S, Chen K, Liu Y, Li H, Ma Y, Zhang Z (2018) FveRGA1, encoding a DELLA protein, negatively regulates runner production in Fragaria vesca. Planta 247: 941–951

Liang Y, Ward S, Li P, Bennett T, Leyser O (2016) SMAX1-LIKE7 Signals from the Nucleus to Regulate Shoot Development in Arabidopsis via Partially EAR Motif-Independent Mechanisms. Plant Cell 28: 1581–1601

Liao X, Li M, Liu B, Yan M, Yu X, Zi H, Liu R, Yamamuro C (2018) Interlinked regulatory loops of ABA catabolism and biosynthesis coordinate fruit growth and ripening in woodland strawberry. Proc Natl Acad Sci U S A 115: e11542-e11550

Liu J, Cheng X, Liu P, Sun J (2017) miR156-Targeted SBP-Box Transcription Factors Interact with DWARF53 to Regulate TEOSINTE BRANCHED1 and BARREN STALK1 Expression in Bread Wheat. Plant Physiol 174: 1931–1948

Liu X, Cheng L, Li R, Cai Y, Wang X, Fu X, Dong X, Qi M, Jiang CZ, Xu T, Li T (2022) The HD-Zip transcription factor SlHB15A regulates abscission by modulating jasmonoyl-isoleucine biosynthesis. Plant Physiol 4: 4

Livak KJ, Schmittgen TD (2001) Analysis of Relative Gene Expression Data Using Real-Time Quantitative PCR and the 2^−ΔΔCT^ Method. Methods 25: 402–408

Ma H, Duan J, Ke J, He Y, Gu X, Xu TH, Yu H, Wang Y, Brunzelle JS, Jiang Y, Rothbart SB, Xu H.E., Li J, Melcher K (2017) A D53 repression motif induces oligomerization of TOPLESS corepressors and promotes assembly of a corepressor-nucleosome complex. Sci. Adv 3: e1601217

Mason MG, Ross JJ, Babst BA, Wienclaw BN, Beveridge CA (2014) Sugar demand, not auxin, is the initial regulator of apical dominance. Proc Natl Acad Sci U S A 111: 6092–6097

Patil SB, Barbier FF, Zhao J, Zafar SA, Uzair M, Sun Y, Fang J, Perez-Garcia MD, Bertheloot J, Sakr S, Fichtner F, Chabikwa TG, Yuan S, Beveridge CA, Li X (2021) Sucrose promotes D53 accumulation and tillering in rice. New Phytol 234: 122–136

Ré DA, Dezar CA, Chan RL, Baldwin IT, Bonaventure G (2010) Nicotiana attenuata NaHD20 plays a role in leaf ABA accumulation during water stress, benzylacetone emission from flowers, and the timing of bolting and flower transitions. J Exp Bot 62: 155–166

Salam BB, Barbier F, Danieli R, Teper-Bamnolker P, Ziv C, Spichal L, Aruchamy K, Shnaider Y, Leibman D, Shaya F, Carmeli-Weissberg M, Gal-On A, Jiang J, Ori N, Beveridge C, Eshel D (2021) Sucrose promotes stem branching through cytokinin. Plant Physiol 185: 1708–1721

Scheres B, Shinohara N, Taylor C, Leyser O (2013) Strigolactone Can Promote or Inhibit Shoot Branching by Triggering Rapid Depletion of the Auxin Efflux Protein PIN1 from the Plasma Membrane. PLOS Biol 11: e1001474

Shabek N, Ticchiarelli F, Mao H, Hinds TR, Leyser O, Zheng N (2018) Structural plasticity of D3-D14 ubiquitin ligase in strigolactone signalling. Nature 563: 652–656

Shao J, Haider I, Xiong L, Zhu X, Hussain RMF, Övernäs E, Meijer AH, Zhang G, Wang M, Bouwmeester HJ, Ouwerkerk PBF (2018) Functional analysis of the HD-Zip transcription factor genes Oshox12 and Oshox14 in rice. Plos One 13: e0199248

Simons JL, Napoli CA, Janssen BJ, Plummer KM, Snowden KC (2007) Analysis of the DECREASED APICAL DOMINANCE genes of petunia in the control of axillary branching. Plant Physiol 143: 697–706

Smith SM, Li J (2014) Signalling and responses to strigolactones and karrikins. Curr Opin Plant Biol 21: 23–29

Son O, Hur YS, Kim YK, Lee HJ, Kim S, Kim MR, Nam KH, Lee MS, Kim BY, Park J, Park J, Lee SC, Hanada A, Yamaguchi S, Lee IJ, Kim SK, Yun DJ, Soderman E, Cheon CI (2010) ATHB12, an ABA-inducible homeodomain-leucine zipper (HD-Zip) protein of Arabidopsis, negatively regulates the growth of the inflorescence stem by decreasing the expression of a gibberellin 20-oxidase gene. Plant Cell Physiol 51: 1537–1547

Song X, Lu Z, Yu H, Shao G, Xiong J, Meng X, Jing Y, Liu G, Xiong G, Duan J, Yao X-F, Liu C-M, Li H, Wang Y, Li J (2017) IPA1 functions as a downstream transcription factor repressed by D53 in strigolactone signaling in rice. Cell Res 27: 1128–1141

Soundappan I, Bennett T, Morffy N, Liang Y, Stanga JP, Abbas A, Leyser O, Nelson DC (2015) SMAX1-LIKE/D53 Family Members Enable Distinct MAX2-Dependent Responses to Strigolactones and Karrikins in Arabidopsis. Plant Cell 27: 3143–3159

Sun H, Guo X, Qi X, Feng F, Xie X, Zhang Y, Zhao Q (2021) SPL14/17 act downstream of strigolactone signalling to modulate rice root elongation in response to nitrate supply. Plant J 106: 649–660

Sun H, Guo X, Zhu X, Gu P, Zhang W, Tao W, Wang D, Wu Y, Zhao Q, Xu G, Fu X, Zhang Y (2023) Strigolactone and gibberellin signaling coordinately regulate metabolic adaptations to changes in nitrogen availability in rice. Mol Plant 16: 588–598

Umehara M, Cao M, Akiyama K, Akatsu T, Seto Y, Hanada A, Li W, Takeda-Kamiya N, Morimoto Y, Yamaguchi S (2015) Structural Requirements of Strigolactones for Shoot Branching Inhibition in Rice and Arabidopsis. Plant Cell Physiol 56: 1059–1072

Valdes AE, Overnas E, Johansson H, Rada-Iglesias A, Engstrom P (2012) The homeodomain-leucine zipper (HD-Zip) class I transcription factors ATHB7 and ATHB12 modulate abscisic acid signalling by regulating protein phosphatase 2C and abscisic acid receptor gene activities. Plant Mol Biol 80: 405–418

van Rongen M, Bennett T, Ticchiarelli F, Leyser O (2019) Connective auxin transport contributes to strigolactone-mediated shoot branching control independent of the transcription factor BRC1. PLoS Genet 15: e1008023

Vegh A, Incze N, Fabian A, Huo H, Bradford KJ, Balazs E, Soos V (2017) Comprehensive analysis of DWARF14-LIKE2 (DLK2) reveals its functional divergence from strigolactone-related paralogs. Front Plant Sci 8: 1641

Wang F, Han T, Song Q, Ye W, Song X, Chu J, Li J, Chen ZJ (2020) The Rice Circadian Clock Regulates Tiller Growth and Panicle Development Through Strigolactone Signaling and Sugar Sensing. Plant Cell 32: 3124–3138

Wang L, Wang B, Jiang L, Liu X, Li X, Lu Z, Meng X, Wang Y, Smith SM, Li J (2015) Strigolactone Signaling in Arabidopsis Regulates Shoot Development by Targeting D53-Like SMXL Repressor Proteins for Ubiquitination and Degradation. Plant Cell 27: 3128–3142

Wang L, Wang B, Yu H, Guo H, Lin T, Kou L, Wang A, Shao N, Ma H, Xiong G, Li X, Yang J, Chu J, Li J (2020) Transcriptional regulation of strigolactone signalling in Arabidopsis. Nature 583: 277–281

Wang M, Pérez-Garcia M-D, Davière J-M, Barbier F, Ogé L, Gentilhomme J, Voisine L, Péron T, Launay-Avon A, Clément G, Baumberger N, Balzergue S, Macherel D, Grappin P, Bertheloot J, Achard P, Hamama L, Sakr S, Cubas P (2021) Outgrowth of the axillary bud in rose is controlled by sugar metabolism and signalling. J Exp Bot 72: 3044–3060

Wang X, Jin B, Yan W, Wang J, Xu J, Cai C, Qi X, Xu Q, Yang X, Xu X, Chen X (2023) Cucumber abscisic acid 8′-hydroxylase Csyf2 regulates yellow flesh by modulating carotenoid biosynthesis. Plant Physiol 193: 1001–1015

Waters MT, Gutjahr C, Bennett T, Nelson DC (2017) Strigolactone Signaling and Evolution. Annu Rev Plant Biol 68: 291–322

Waters MT, Nelson DC, Scaffidi A, Flematti GR, Sun YK, Dixon KW, Smith SM (2012) Specialisation within the DWARF14 protein family confers distinct responses to karrikins and strigolactones in Arabidopsis. Development 139: 1285–1295

Wu H, Li H, Chen H, Qi Q, Ding Q, Xue J, Ding J, Jiang X, Hou X, Li Y (2019) Identification and expression analysis of strigolactone biosynthetic and signaling genes reveal strigolactones are involved in fruit development of the woodland strawberry (Fragaria vesca). BMC Plant Biol 19: 1–19

Wu K, Wang S, Song W, Zhang J, Wang Y, Liu Q, Yu J, Ye Y, Li S, Chen J, Zhao Y, Wang J, Wu X, Wang M, Zhang Y, Liu B, Wu Y, Harberd NP, Fu X (2020) Enhanced sustainable green revolution yield via nitrogen-responsive chromatin modulation in rice. Science 367: eaaz2046

Xie Y, Liu Y, Ma M, Zhou Q, Zhao Y, Zhao B, Wang B, Wei H, Wang H (2020) Arabidopsis FHY3 and FAR1 integrate light and strigolactone signaling to regulate branching. Nat Commun 11: 1955

Yao R, Ming Z, Yan L, Li S, Wang F, Ma S, Yu C, Yang M, Chen L, Chen L, Li Y, Yan C, Miao D, Sun Z, Yan J, Sun Y, Wang L, Chu J, Fan S, He W, Deng H, Nan F, Li J, Rao Z, Lou Z, Xie D (2016) DWARF14 is a non-canonical hormone receptor for strigolactone. Nature 536: 469–473

Yao R, Wang L, Li Y, Chen L, Li S, Du X, Wang B, Yan J, Li J, Xie D (2018) Rice DWARF14 acts as an unconventional hormone receptor for strigolactone. J Exp Bot 69: 2355–2365

Zhang J, Mazur E, Balla J, Gallei M, Kalousek P, Medveďová Z, Li Y, Wang Y, Prát T, Vasileva M, Reinöhl V, Procházka S, Halouzka R, Tarkowski P, Luschnig C, Brewer PB, Friml J (2020) Strigolactones inhibit auxin feedback on PIN-dependent auxin transport canalization. Nat Commun 11: 3508

Zhang Y, Zhang Y, Sun Q, Lu S, Chai L, Ye J, Deng X (2021) Citrus transcription factor CsHB5 regulates abscisic acid biosynthetic genes and promotes senescence. Plant J 108: 151–168

Zhao H, Wang Y, Zhao S, Fu Y, Zhu L (2021) HOMEOBOX PROTEIN 24 mediates the conversion of indole-3-butyric acid to indole-3-acetic acid to promote root hair elongation. New Phytol 232: 2057–2070

Zhao S, Wang H, Jia X, Gao H, Mao K, Ma F (2021) The HD-Zip I transcription factor MdHB7-like confers tolerance to salinity in transgenic apple (Malus domestica). Physiol Plant 172: 1452–1464

Zhou F, Lin Q, Zhu L, Ren Y, Zhou K, Shabek N, Wu F, Mao H, Dong W, Gan L, Ma W, Gao H, Chen J, Yang C, Wang D, Tan J, Zhang X, Guo X, Wang J, Jiang L, Liu X, Chen W, Chu J, Yan C, Ueno K, Ito S, Asami T, Cheng Z, Wang J, Lei C, Zhai H, Wu C, Wang H, Zheng N, Wan J (2013) D14–SCF^D3^-dependent degradation of D53 regulates strigolactone signalling. Nature 504: 406–410

